# Functional genetics reveals a role for the proteasome in regulating centromere assembly

**DOI:** 10.64898/2026.10.14.759079

**Authors:** Océane Marescal, Kuan-Chung Su, Matteo Di Bernardo, Kaitlyn Manzer, Jiayi He, Roshan Kern, Iain Cheeseman

## Abstract

Centromeres link chromosomes to spindle microtubules to direct their segregation during mitosis. The vertebrate centromere is built on two critical DNA-binding proteins, the histone H3-variant CENP-A and histone fold protein CENP-T. CENP-T is a potent scaffold for kinetochore assembly, such that concentrating it at ectopic sites is sufficient to build kinetochore-like structures. How cells prevent excess CENP-T from acting at non-centromere sites remains unclear. Here, using optical pooled screening to systematically define the requirements for CENP-A and CENP-T localization, we recover the established CENP-A deposition machinery and reveal a specific role for the proteasome in controlling CENP-T levels. We find that non-centromere CENP-T is degraded in a ubiquitin- and proteasome-dependent manner. Whereas centromere-incorporated CENP-T is stable, CENP-T that is not bound to centromeres is rapidly turned over, with the CENP-T histone fold region sufficient to confer proteasome sensitivity. Quiescent cells, which disassemble their centromeres, continue to express CENP-T but eliminate it through continuous degradation. Thus, whereas CENP-A is controlled primarily through regulated deposition, cells additionally control CENP-T by selectively degrading the unincorporated protein.

## Introduction

The centromere is the region of the chromosome responsible for its segregation during cell division, serving as the point of contact between chromosomal DNA and spindle microtubules [1]. In vertebrates, the centromere is defined epigenetically by the presence of CENP-A, a centromere-specific histone H3 variant. In addition to CENP-A, the centromere is composed of 16 constitutively localized protein components [2–4]: CENP-C, the CENP-T/W/S/X complex, the CENP-H/I/K/M complex, the CENP-L/N complex, and the CENP-O/P/Q/U/R complex. Together, these components make up the Constitutive Centromere Associated Network (CCAN), which recruits the microtubule-binding proteins of the outer kinetochore to the chromosomes during mitosis.

The centromere has two primary DNA binding components: CENP-A and CENP-T (Fig. 1A). CENP-A is the key epigenetic marker for centromere identity; its loss abolishes kinetochore assembly, whereas targeting CENP-A to another chromosomal locus is sufficient to recruit centromere proteins to an ectopic site [5]. To maintain centromere identity, CENP-A is redeposited at centromeres once per cell cycle in early G1 in a tightly regulated manner [6, 7]. The specialized CENP-A deposition machinery includes the chaperone HJURP [8, 9] and components of the Mis18 complex [10]. However, CENP-A alone is not sufficient for kinetochore assembly, requiring other components such as CENP-C and CENP-T for full kinetochore formation [11–13]. CENP-T serves to recruit the outer kinetochore during cell division. Its N-terminus binds to microtubule-binding components, NDC80 and MIS12 [14, 15], while its C-terminal histone fold region binds to the DNA [16], thus allowing CENP-T to act as a link between the chromosome and the microtubules. CENP-T levels must be tightly controlled. Indeed, artificially creating a high local concentration of CENP-T results in the formation of ectopic kinetochore-like particles and the potent recruitment of outer kinetochore proteins, resulting in mitotic defects [17]. Thus, CENP-A and CENP-T play essential roles in centromere identity and function, and understanding what regulates their stability, incorporation, and levels is crucial.

**Figure 1:**
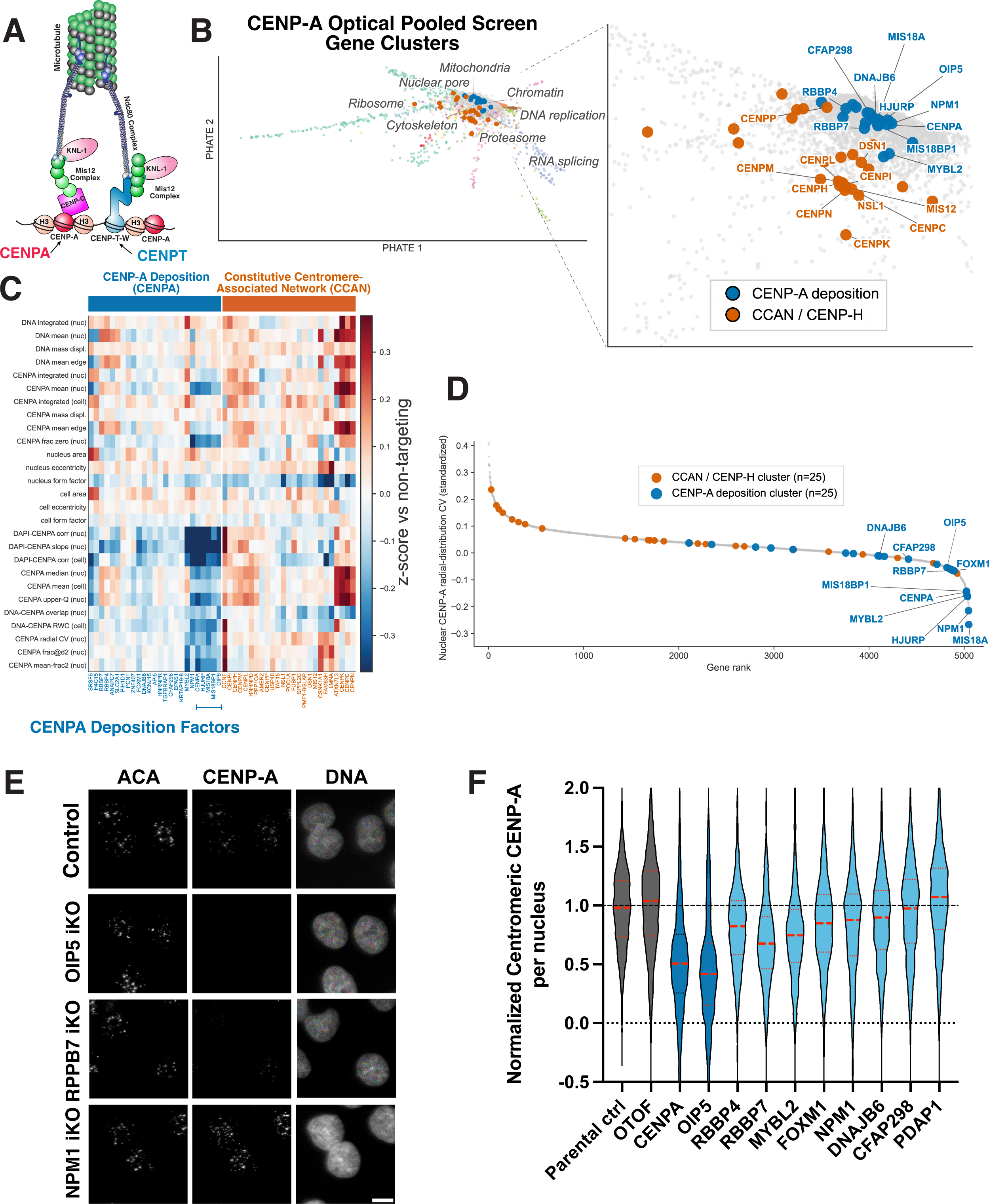
Optical Pooled Screening identifies molecular requirements for CENP-A localization. A. Schematic of the kinetochore, highlighting the centromere DNA binding proteins CENP-A and CENP-T. B. PHATE plot showing the phenotypic distribution of gene knockouts from the Optical Pooled Screening (OPS) analysis of CENP-A. Right, inset showing blow up of the clusters of gene knockouts for centromere biology and CENP-A deposition. C. Heatmap showing representative features obtained from the OPS analysis for the centromere and CENP-A deposition clusters. D. Waterfall plot showing the relative nuclear CENP-A radial distribution obtained from the OPS analysis, which is substantially decreased in gene knockouts for established CENPA deposition factors. E. Representative immunofluorescence images showing the localization of anti-centromere antibodies (ACA) and CENP-A and DNA in individual knockout cell lines. Scale bar, 10 µm. F. Graph showing the quantification of normalized centromere CENP-A intensity in individual cells from E. n>2000 cells per condition from 3 independent experiments

Here, we conduct Optical Pool Screens (OPS) to identify regulators of CENP-A and CENP-T biology. OPS combines CRISPR-Cas9 mediated inducible gene knockouts with microscopy-based readouts for cellular phenotypes. Previous OPS analyses typically use broadly relevant cellular markers that report on the functions of multiple genes [18, 19], analogous to the concept of cell painting [20]. However, OPS is also structured to allow a systematic analysis for the requirements of any cell biological process that is compatible with microscopy-based imaging. Here, we leverage this strategy to analyze the requirements for centromere assembly and regulation. By analyzing CENP-A localization systematically across all human fitness-conferring genes, our CENP-A screen effectively identified the established deposition factors for CENP-A, as well as identifying chromatin-associated factors that influence CENP-A distribution. By analyzing CENP-T levels and chromatin recruitment, our large-scale analysis of CENP-T localization revealed a key role for proteasomal degradation in regulating CENP-T homeostasis. We find that CENP-T is tightly controlled in a proteasome- and ubiquitin-dependent manner and that this degradation serves to prevent the accumulation of excess non-centromere bound CENP-T in the nucleus. Finally, we show that CENP-T degradation occurs constitutively in quiescent cells to ensure that CENP-T is eliminated from this non-dividing cell state.

## Results and Discussion

### Optical Pooled Screening systematically identifies CENP-A assembly factors

To define the requirements for centromere assembly and regulation in human HeLa cells, we began by conducting optical pooled screening (OPS) using a Cas9 sgRNA library targeting all fitness-conferring genes [18] and immuno-staining for the centromere-specific histone H3 variant, CENP-A (Fig. 1A). Following knockout induction for 78 hours, we fixed cells and conducted phenotypic imaging and *in situ* sequencing. We then conducted an in-depth computational analysis of the image data using the Brieflow pipeline [21], including the extraction of image features based on Cell Profiler ([22]; Supplemental Table 1). Based the combination of these features, we were able to cluster groups of genes with related morphological phenotypes in PHATE space (Fig. 1B; Supplemental Table 1). Based on features derived from just DNA and CENP-A staining, we were able to group genes into major functional categories, including the ribosome, RNA splicing machinery, mitochondria, and others (Fig. 1B). This analysis shared features also revealed two clear gene clusters with relevance to CENP-A biology (Fig. 1B, C). The first cluster contained established centromere and kinetochore proteins, including components of the Constitutive Centromere-Associated Network (CCAN), whereas the second contained each of the established CENP-A deposition factors (CENPA itself, HJURP, MIS18BP1, OIP5, and MIS18A).

Analysis of the morphological features from the CENP-A-containing gene cluster indicated a substantial decrease in nuclear CENPA levels (Fig. 1D), with a particular reduction in the nuclear DAPI-CENPA correlation and the nuclear DAPI-CENPA slope for established CENPA deposition factors. To test the contributions of other genes in this cluster, we created individual inducible knockout cell lines [23, 24]. Analysis of normalized centromere CENPA intensity confirmed a substantial decrease in CENPA levels in knockouts for CENPA and the CENPA deposition factor OIP5. In addition, we observed modestly decreased centromere CENPA levels in knockouts of the chromatin remodeling factors RBBP4 and RBBP7 and the transcription factor MYBL2 (Fig. 1E,F), consistent with prior work [25–27]. In total, this analysis demonstrates the ability of large-scale optical pooled screening to identify gene targets with specific roles in centromere biology.

### Optical Pooled Screening identifies players in CENP-T biology

To uncover novel players and mechanisms in centromere biology, we next leveraged our OPS pipeline in HeLa cells to visualize additional centromere proteins using Anti-Centromere Antibodies (ACA; which primarily detect the alpha-satellite binding protein CENPB) and antibodies against the histone-fold containing protein CENP-T [13]. Clustering analysis of the complete phenotypic features from this screen identified a clear gene cluster containing CENP-T together with other centromere proteins, including the obligate CENP-T binding partner CENP-W, as well as CENP-C, CENP-H, CENP-I, CENP-K, CENP-L, CENP-M, and CENP-N (Fig. 2A; Supplemental Fig. 1A; Supplemental Table 1). Strikingly, this analysis also resulted in distinct clusters of proteasome subunits, which were not visible in the CENPA screen.

**Figure 2:**
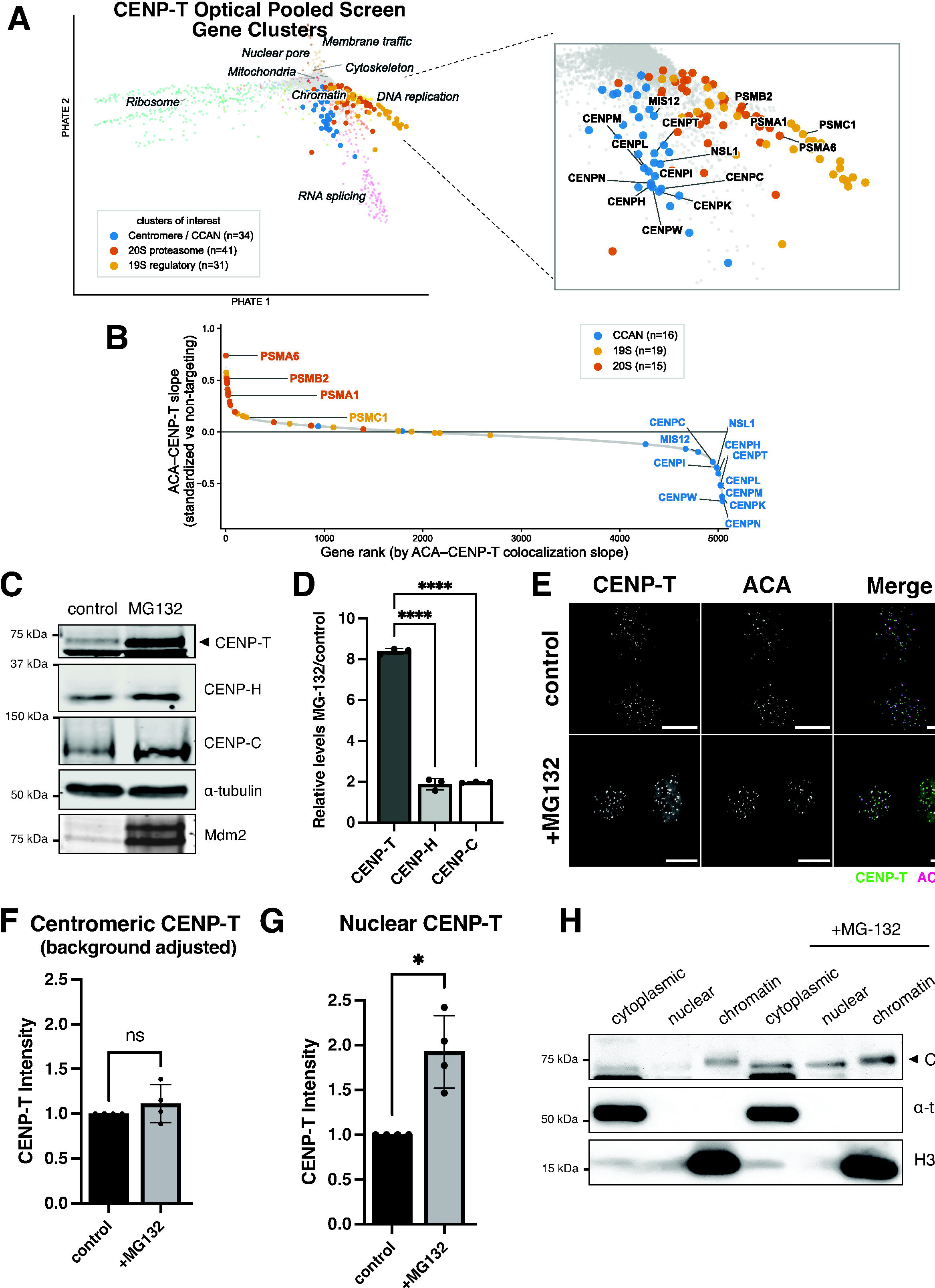
CENP-T is degraded in a proteasome-dependent manner. A. PHATE plot showing the phenotypic distribution of gene knockouts from the Optical Pooled Screening (OPS) analysis of ACA and CENP-T. Right, inset showing blow up of the clusters of gene knockouts for centromere biology and the 19S and 20S proteasome subunits. B. Waterfall plot for the relative ACA vs. CENP-T co-localization slope highlight gene targets that reduce CENP-T levels (CCAN proteins, blue) or increase CENP-T levels (proteasome subunits, yellow and orange). C. Western blot of control cells and cells treated for 24 hours with 10 µM MG-132. Three separate blots are shown. The first blot was incubated with anti-CENP-T and anti-CENP-H antibodies. The second blot was incubated with anti-CENP-C antibody. And the third with anti-MDM2 antibodies. MDM2 is known to be degraded rapidly and shows efficacy of MG-132 treatment. α-tubulin is used as a loading control and is shown for the CENP-T/CENP-H blot, the same amounts were loaded for all three blots. D. Graph of the relative levels of centromere proteins in MG-132-treated vs. control cells by quantitative western blot analysis. Three separate biological replicates were run and bands were quantified. Centromere protein band intensities were divided by loading control band intensities. Values for MG-132 samples were divided by control sample values. Bars represent mean ± standard deviation. **** represents p<0.0001. E. Representative immunofluorescence images of control cells and cells treated for 24 hours with 10 µM MG-132. Cells were stained with CENP-T antibody and anti-centromere antibodies (ACA). Scale bar, 10 µm. F. Graph showing the CENP-T intensity measured at the centromeres minus background intensity of control cells vs. cells treated with MG-132 from Figure 2E. Graph was generated from 4 separate biological replicates. Bars represent mean ± standard deviation. ns = not significant. Values normalized to control. G. Graph showing the CENP-T intensity measured at the nucleus of control cells vs. cells treated with MG-132 from Figure 2E. Graph was generated from 4 separate biological replicates. Bars represent mean ± standard deviation. * represents p = 0.0196. Values normalized to control. H. Western blot of nuclear fractionation experiment. Cytoplasmic, nuclear, and chromatin fractions are indicated, with or without addition of 10 µM MG-132. Blot was incubated with CENP-T antibody. α-tubulin is a control for cytoplasmic fraction, H3 is a control for chromatin-bound.

We next evaluated changes in CENP-T abundance. To do so, we examined mean nuclear CENP-T intensity for each gene knockout against mean nuclear ACA intensity (Figure 2B), allowing us to visualize gene knockouts that specifically led to increased or decreased CENP-T intensity. As expected, knockout of either CENP-T or its binding partner CENP-W led to decreased CENP-T nuclear intensity without affecting ACA levels (Fig. 2B), validating this approach. In addition, we found that gene knockouts for multiple subunits of the 20S proteasome led to highly increased CENP-T intensity relative to ACA levels (Fig. 2B). In contrast to this potent effect on CENP-T levels, in our OPS analysis of CENP-A localization, we did not observe a clear change in CENP-A levels in proteasome knockouts (Fig. 1; Supplemental Table 1). Thus, this change in CENP-T levels appears to be specific to CENP-T and not a general increase in protein levels. Together, this analysis suggests that CENP-T levels may be controlled by protein degradation.

### CENP-T is degraded in a proteasome-dependent manner

The regulation of CENP-T is critical for ensuring proper chromosome segregation. Indeed, an increased local concentration of CENP-T is sufficient to induce the formation of ectopic kinetochore-like particles [17] such that restraining the levels of CENP-T at non-centromere regions is of critical importance. We found that Cas9-based knockouts for multiple different proteasome subunits resulted in increased CENP-T based on optical pooled screening (Fig. 2B, Supplemental Fig. 1A). Therefore, we next investigated whether proteasome activity could play a role in regulating CENP-T levels. Treatment of cells for 24 hours with MG-132, a chemical inhibitor of proteasome activity, led to ∼8-fold increase in CENP-T protein levels based on western blot analysis, whereas the levels of the CCAN proteins CENP-H and CENP-C remained relatively stable (Fig. 2C, D). Thus, CENP-T, but not other centromere proteins, accumulates upon proteasome inhibition. Similarly, treatment of cells for 24 hours with TAK-243, which prevents ubiquitination of proteins by inhibiting the E1 activating enzyme, also led to CENP-T protein accumulation (Supplemental Fig. 1B). Thus, under steady state, CENP-T protein is degraded in a proteasome- and ubiquitin-dependent manner.

We next sought to determine the behavior of stabilized CENP-T protein following proteasome inhibition. Although cells treated with MG-132 did not have increased CENP-T localization to centromeres based on immunofluorescence (Fig. 2E, F), we observed an accumulation in nuclear CENP-T intensity (Fig. 2E, G). This increase in nuclear CENP-T could reflect either increased chromatin-bound CENP-T or increased soluble nuclear-localized CENP-T that is not bound to chromatin. Thus, we additionally conducted chromatin fractionation to distinguish these two populations. Although addition of MG-132 led to only a small increase in the amount of chromatin-bound CENP-T, we found that the levels of unbound nuclear CENP-T in MG-132-treated cells were much higher (Fig. 2H). Thus, proteasome inhibition results in CENP-T accumulation, with an increased nucleoplasmic pool that does not bind stably to chromatin or centromeres.

### Non-chromatin-bound CENP-T is targeted for degradation

As nuclear CENP-T accumulated upon proteasome inhibition, we next sought to evaluate the contribution of CENP-T degradation. To test this, we conducted a cycloheximide chase assay in which we inhibited new protein translation using cycloheximide and monitored the levels of CENP-T over time. Surprisingly, addition of cycloheximide led to little change in overall CENP-T protein levels or centromere intensity (Fig. 3A, B), indicating that there is minimal ongoing degradation of centromere-localized CENP-T protein under normal circumstances. Thus, we next induced the accumulation of excess non-centromeric CENP-T using MG-132 treatment, and then measured CENP-T turnover following MG-132 washout and cycloheximide treatment (Fig. 3C). In contrast to CENP-T in untreated cells (Fig. 3A), the excess pool of nuclear CENP-T was quickly reduced within 8 hours following cycloheximide addition (Fig. 3D), indicating that it is rapidly turned over.

**Figure 3:**
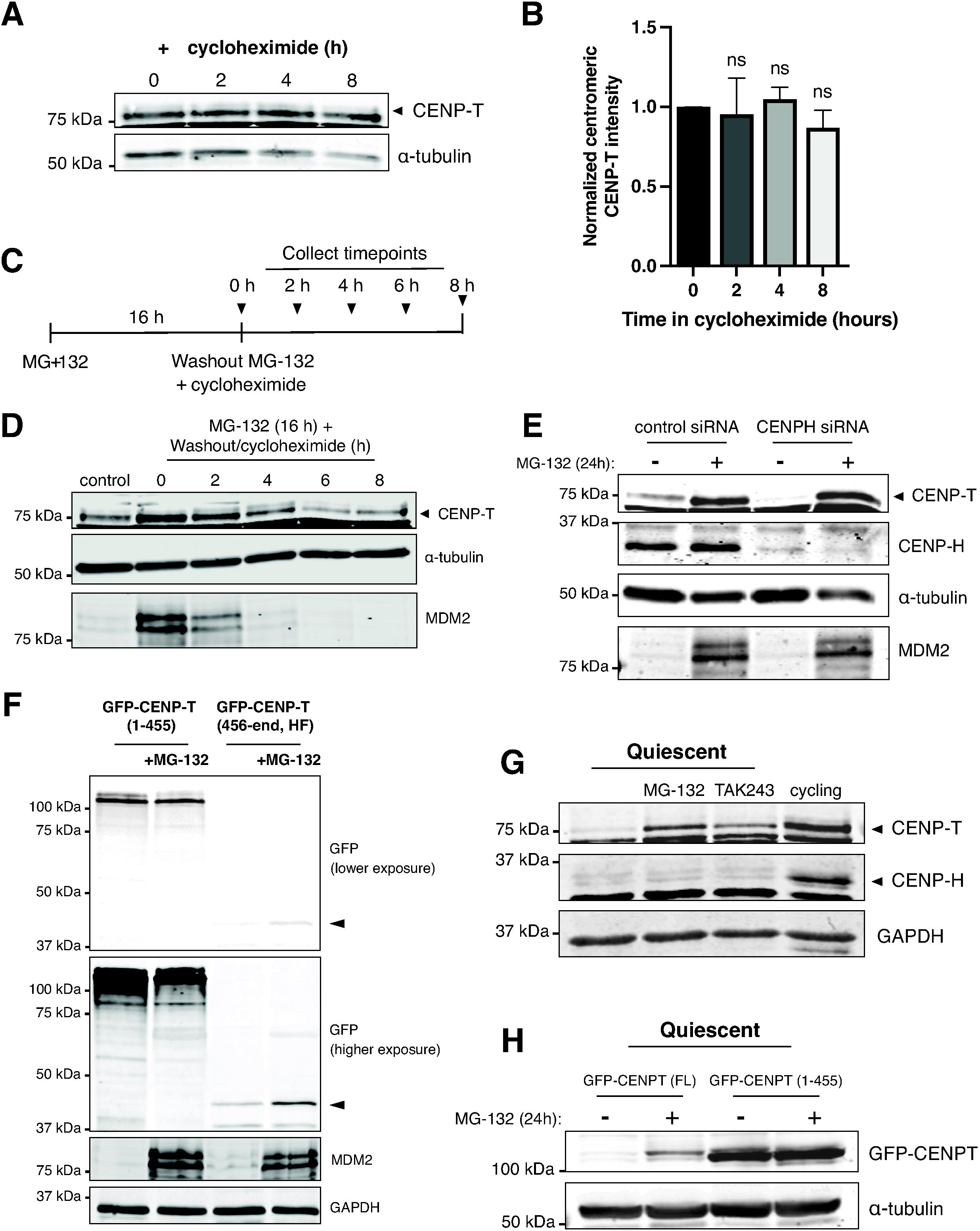
Non-chromatin-bound CENP-T is targeted for degradation. A. Western blot of cells treated with the indicated amount of time of cycloheximide (50 µg/mL). Blot was incubated with anti-CENP-T antibody. α-tubulin serves as a loading control. B. Graph showing centromeric CENP-T intensity over time of cycloheximide treatment quantified from immunofluorescence experiments. Three separate biological replicates/experiments were quantified. ns = not significant. Values normalized to 0-hour time point. n = 6179, 6987, 5139, and 4415 centromeres quantified for 0-, 2-, 4-, and 8-hour time points, respectively. C. Diagram showing experimental set-up for experiment shown in Figure 3D. Cells were treated with MG-132 for 16 hours. MG-132 was then washed out and cycloheximide was added for the indicated amounts of time before collection. D. Western blot showing results from the experiment diagrammed in Figure 3C. Blot was incubated in anti-CENP-T antibodies. A separate blot was incubated in anti-MDM2 antibody to show efficacy of MG-132 treatment. α-tubulin serves as a loading control. E. Western blot of cells treated with 50 nM control siRNAs or 50 nM CENPH siRNAs for 72 hours, with or without the addition of 10 µM MG-132 for 24 hours. Blot was incubated with anti-CENP-T and anti-CENP-H antibodies. MDM2 controls for MG-132 efficacy and α-tubulin serves as a loading control. F. Western blot of truncated GFP-CENP-T cell lines with or without 24 hours of MG-132 treatment (10 µM). Blot was incubated with anti-GFP antibodies. MDM2 controls for MG-132 efficacy and GAPDH serves as a loading control. Lower and higher exposures are shown. Arrow indicates the band for the GFP-CENPT (456-end) construct. G. Western blot showing quiescent cells with no drug, 24 hours of 10 µM MG-132, 24 hours of 400 nM TAK-243, or cycling cells. Blot was incubated with CENP-T and CENP-H antibodies. GAPDH is used as a loading control. Cells were in quiescence for 7 days. H. Blot showing quiescent cell lines with GFP-CENPT constructs: Full length (FL) and N-terminal CENPT (1-455), with or without 24 hours of 10 µM MG-132 treatment. Blot was incubated in anti-GFP antibody. α-tubulin serves as a loading control. Cells were in quiescence for 7 days.

We next evaluated whether proteasomal CENP-T degradation represents a mechanism for downregulating excess CENP-T to maintain normal homeostatic protein levels or whether degradation occurs specifically for unincorporated CENP-T. To test whether creating a pool of non-centromere bound CENP-T without increasing total CENP-T levels results in its degradation, we took advantage of the interdependent nature of centromere protein localization. Because CENP-T localization to centromeres requires the presence of CENP-H [24], we used siRNAs to deplete CENP-H and induce CENP-T delocalization. Delocalized CENP-T protein that was evicted from centromeres was lost in CENP-H knockdown cells based on western blots (Fig. 3E). However, CENP-T levels were rescued in CENP-H depleted cells by proteasome inhibition (Fig. 3E). Thus, unincorporated CENP-T is rapidly degraded in a proteasome-dependent manner regardless of the total CENP-T levels.

Finally, we sought to determine the region of CENP-T responsible for its degradation. CENP-T is composed of two regions: a disordered N-terminal domain with multiple binding sites for outer kinetochore proteins [13, 14] and a C-terminal histone fold domain, which associates with CENP-W to incorporate the CENP-T-W dimer into the chromatin [16]. Given that centromere-bound CENP-T was stable while unincorporated CENP-T was degraded, we reasoned that exposure of the CENP-T histone fold in non-localized CENP-T could trigger its degradation. To test this, we created two constructs containing different truncated domains of CENP-T as GFP fusions (Supplemental Fig. 2A). Although the levels of the N-terminal domain of CENP-T (GFP-CENP-T, residues 1-455) were not affected by MG-132 treatment, the C-terminal histone fold region of CENP-T (GFP-CENP-T, residues 456-end) accumulated following MG-132 addition (Fig. 3F; Supplemental Fig. 2B), indicating that this domain alone can induce the degradation of the GFP fusion.

Together, these data indicate that non-centromere localized, unbound CENP-T is degraded by the proteasome and that exposure of the histone fold could act as a signal to target the protein for degradation.

### Quiescent cells degrade CENP-T

We next considered whether there are physiological contexts in which it is important to regulate CENP-T protein levels. Quiescence is a state of reversible cell cycle arrest [28] in which the centromere is largely disassembled, resulting in the delocalization of the CCAN proteins including CENP-T and CENP-H [29, 30]. Our previous work found that quiescent cells transcriptionally downregulate the levels of most centromere protein genes, but that CENP-T mRNA expression remains high [30]. Despite this, quiescent cells lack detectable CENP-T protein [29, 30]. We therefore considered whether quiescent cells regulate CENP-T post-transcriptionally via proteasomal degradation. To test this, we treated quiescent cells with either MG-132 or TAK-243 to inhibit proteasome activity and ubiquitination, respectively. Whereas untreated quiescent cells did not have detectable CENP-T protein, both MG-132 and TAK-243 treatment led to the rapid recovery of CENP-T protein levels within only 4 hours of proteasome inhibition (Fig. 3G; Supplemental Fig. 2C, D). Conversely, CENP-H protein remained low even in MG-132 treated cells (Fig. 3G), consistent with our previous work showing that the downregulation of CENP-H during quiescence occurs at the transcriptional level [30]. However, stabilized CENP-T protein did not regain centromere localization (Supplemental Fig. 2E, F), as expected due to the absence of other centromere proteins. These results suggest that cells continuously produce and degrade CENP-T during quiescence. Indeed, inhibition of new CENP-T translation by cycloheximide treatment prevented recovery of CENP-T protein levels upon proteasome inhibition (Supplemental Fig. 2D). Thus, quiescent cells use proteasome-mediated degradation to post-transcriptionally regulate CENP-T protein levels and centromere disassembly during quiescence, which may enable the rapid production of CENP-T following the exit from quiescence.

### Optical Pooled Screening provides novel insights into centromere biology

The results described here highlight the ability of large-scale microscopy-based analysis enabled by optical pooled screening to reveal new insights into targeted cell biological questions, including centromere regulation. Regulating the concentration of CENP-T is essential for proper centromere formation and function. As excess amounts of CENP-T could lead to the formation of ectopic kinetochores and result in chromosome mis-segregation [17], the ability to eliminate excess non-centromeric CENP-T is crucial. Here, we find that cells degrade unbound CENP-T in a proteasome-dependent manner to maintain proper levels of CENP-T protein. We find that this degradation is mediated by the histone fold region of the protein, which is occluded when CENP-T is appropriately incorporated with DNA at centromeres, but exposed in unbound CENP-T.

Centromere proteins play critical roles in the segregation of chromosomes during cell division [1]. As quiescent cells reside in a non-dividing state, they do not require a centromere and therefore dismantle it during quiescence entry [30]. Whereas the downregulation of other centromere proteins during quiescence occurs at the transcriptional level [30], we find that quiescent cells continue to produce CENP-T protein, but eliminate it through continuous protein degradation. Quiescent cells are characterized by their ability to rapidly reenter the cell cycle following stimulation [28]. The CENP-T degradation mechanism may ensure that no CENP-T protein remains in a quiescent cell, while also allowing for a rapid increase in CENP-T levels upon the resumption of cell division. Thus, continuous production of CENP-T could keep quiescent cells poised to rapidly reassemble its centromere at any moment upon receiving a signal to return to growth. Together, these studies reveal new insights into centromere regulation and provide a model for leveraging optical pooled screening to define new cell biological principles.

## Acknowledgements

The authors thank the members of the Cheeseman laboratory for their advice and support. The authors would like to thank the Whitehead Flow Cytometry Core for cell sorting and analysis. This work was supported by grants from NIGMS (R35GM126930), the Chan Zuckerberg Initiative (Cell Biology @ Scale), and an MIT (Bose Fellowship) to IMC, and a fellowship to MD from the National Science Foundation.

## Materials and Methods

### Cell culture and reagents

HeLa and RPE1 cells were grown in Dulbecco’s modified Eagle medium (DMEM) supplemented with 10% or 0% (for quiescence) fetal bovine serum (FBS), 100 U/ml penicillin and streptomycin, and 2 mM L-glutamine at 37°C with 5% CO_2_. For quiescence, RPE1 cells were allowed to grow until confluence and left to contact inhibit for 1-2 days. Afterwards, media was removed from cells and replaced with 0% FBS serum starvation media. Serum starvation media was replaced every day for 7 days. Drugs used on cell lines were: MG-132 (10 mM, Enzo Life Sciences) and cycloheximide (50 µg/ml; Sigma-Aldrich). Doxycycline-inducible cell lines were cultured in medium containing tetracycline-free FBS, and were induced with 1 µg/mL doxycycline hyclate. Doxycycline-containing media was renewed every ∼24 hours. Cells were routinely monitored for mycoplasma contamination using commercial detection kits.

### Cell Line Generation

Cell lines used in this study are listed in Table 1. Inducible knock out cell lines targeting individual genes derived by lentiviral transduction of sgRNA into HeLa cells expressing inducible cas9 (cTT20). sgRNA constructs cell lines were generated by primer annealing and ligation into sgOPTI: Addgene #85681 [24, 31]. A control sgRNA with a single target site within the non-essential LBR gene (HS1) [32] was used for comparison. Lentivirus was produced by transfection of HEK-293 cells with sgRNA-plasmid, pCMV-VSV-G, psPAX2. Transduced cells were selected with 0.5 μg/ml puromycin to generate polyclonal cell lines.

GFP-CENP-T cell lines (including GFP-CENP-T (1-455; pOM46), GFP-CENP-T (456-end; pOM47), GFP-CENP-T (pOM11) Full length in HeLa and RPE) were generated by retroviral transduction. Retrovirus was made by transfection of GFP-CENP-T-containing retroviral plasmid and pCMV-VSV-G into HEK-293GP cells. Cells were then selected using 2 μg/ml for HeLa cell lines and 10 μg/ml for RPE cell lines. Blasticidin or sorted for GFP by fluorescence-activated cell sorting.

### Western Blot

For western blot experiments, cells were harvested and frozen, with cell pellets stored at -80°C or immediately lysed on ice for 25 minutes in urea lysis buffer (50 mM Tris pH 7.5, 150 mM NaCl, 0.5% NP-40, 0.1% sodium dodecyl sulfate, 6.5 M Urea, 1X complete EDTA-free protease inhibitor cocktail (Roche), 1 mM phenylmethylsulfonyl fluoride). Cells were then centrifuged for 20 minutes and supernatant was collected. Laemmli sample buffer and 2-mercaptoethanol were added to the lysates, which were then heated at 95 °C for 5 minutes. Samples were loaded on 12% acrylamide gels for SDS-PAGE and transferred to nitrocellulose or PVDK membranes (Cytiva). Membranes were blocked for 30 minutes to 1 hour in 2.5% milk in TBS-T. Primary antibodies (CENP-T: 1 in 2500 [13]; CENP-H: 1:1000 [13]; MDM2: 1:1000; Cell Signaling Technologies 86934S; TubA4A; 1:4000; Sigma; T6199; H3 1:1000; Abcam EPR17899: GAPDH; 1:4000) were added for one hour or overnight, membrane was washed, and then secondary antibodies were added for one hour. Secondary antibodies were either HRP-conjugated or IRDye secondary antibodies for LI-COR imaging. After washing, membranes were incubated in either clarity-enhanced chemiluminescence substrate (Bio-Rad) and imaged using the KwikQuant Imager (Kindle Biosciences) or imaged using the Odyssey CLx Imager (LI-COR).

### Immunofluorescence

Cells were plated on 24 well glass bottom plates (Cellvis) or poly-L-lysine (Sigma Aldrich)-coated or uncoated (for RPE1 cells) coverslips. Cells were then fixed in 4% Formaldehyde diluted in PBS or in PBS with 0.5% TritonX-100 (PBS-Tx). Cells were washed and then blocked in blocking buffer (20 mM Tris-HCl, 150 mM NaCl, 0.1% Triton X-100, 3% bovine serum albumin, 0.1% NaN_3_, pH 7.5) for 30 minutes or overnight at 4°C. Primary antibodies (CENP-A; 1:1000; Invitrogen MA1-20832, CENP-T/W; 2.5mg/L [13]; Vimentin; 1:500 Invitrogen PA1-16759 and Anti-Centromere Protein Antibody 1:100; Antibodiesinc 15-234; TubA4A; 1:4000; Sigma; T6199) were diluted in blocking buffer and added for 1 hour or overnight. Cells were washed and secondary antibodies were added for 1 hour. GFP booster 1:200 (Chromotek, gb2AF488) was applied with secondary antibodies from Jackson ImmunoResearch Laboratories were Cy2-, Cy3-, Alexa 647, or Cy5-conjugated. Then, 1 µg/ml Hoechst-33342 (Sigma Aldrich) was added for 10 minutes. ProLong Gold antifade reagent (Invitrogen) was used to mount coverslips onto slides. Images were taken using a DeltaVision Ultra High-Resolution microscope and analyzed using Fiji/ImageJ. 24 well plates were imaged with PBS using Nikon Ti2 widefield microscope with 21 z-slices at 0.3 μm intervals, max intensity projected and quantified using cell profiler [33].

### Nuclear Fractionation

Cells were harvested and counted to ensure the same number of cells for each condition. Cells were centrifuged at 500xg and cell pellet was resuspended in ice cold Hypotonic Lysis Buffer (HLB; 10 mM Tris pH 7.5, 10 mM NaCl, 3 mM MgCl_2_, 0.3% NP-40, 10% glycerol) at 1 mL HLB per 10 million cells. Roche Protease inhibitor tablet, 0.1M Sodium Orthovanadate, and 0.1M NaF were added. Cells were incubated for 10 minutes and then vortexed. Cells were then centrifuged at 800xg at 4°C for 8 minutes. Supernatant is the cytoplasmic fraction and was collected and 5 M NaCl was added to the cytoplasmic fraction to 140 mM final concentration. Nuclear pellet was washed 4 times with HLB, each time centrifuged at 200xg for 2 minutes. Nuclear pellet was then resuspended in 500 µL Modified Wuarin Schibler Buffer (MWS; 10 mM Tris-HCl pH7, 4 mM EDTA, 0.3 M NaCl, 1M Urea, 1% NP-40), with Roche Protease inhibitor tablet, NaOrthovandate, and NaF and vortexed. This was incubated for 15 minutes, then centrifuged at 1000xg at 4°C for 5 minutes. Supernatant is the nucleoplasmic fraction and pellet is the chromatin-bound fraction. Chromatin fraction was washed 2 times with MWS buffer, with centrifugation at 500xg between each wash for 3 minutes cold. 500 µL of Nuclear Lysis Buffer (NLB; 20 mM Tris pH7.5, 150 mM KCl, 3 mM MgCl_2_, 0.3% NP-40, 10% glycerol) was added to the chromatin pellet, which was then sonicated 3 times at 20% amplitude for 15 seconds. All fractions were then centrifuged at 18,000xg for 15 minutes at 4°C and supernatant was flash frozen in liquid nitrogen at stored at -80°C.

### siRNA treatment

Custom siRNAs against CENPH (5’-GAUCAUACGACAAAACCUA) and a non-targeting control pool (D-001206-13) were obtained from Dharmacon. siRNAs were used at a final concentration of 50 nM. siRNAs and Lipofectamine RNAiMAX (Invitrogen) were first mixed at equal volume in serum-free media (Opti-MEM, ThermoFischer) and vortexed. After 20 minutes of incubation, the mixture was added to cells. Media was changed after 12-24 hours to full-serum media.

### Optical pooled screening

Optical pooled screens were carried out as previously described [18, 34]. The essential-gene inducible KO HeLa cell library was induced with 1 μg/mL doxycycline hyclate for 78 hours and seeded at a density of 350k cells/well in 6-well glass-bottom plates (Cellvis, P06-1.5H-N) 48 hours prior to fixation. Cells were fixed with 4% formaldehyde (Electron Microscopy Sciences, 15710) and 0.007% glutaraldehyde (v/v) in PBS for 25 minutes. All steps used RNase free reagents to preserve mRNA integrity up until the reverse transcription step was complete. The cells were then washed with PBS and permeabilized with 70% ethanol for 30 minutes, followed by two washes in PBS plus 0.05% Tween (PBS-T). Cells were incubated with primary antibodies (CENP-A; 1:1000; Invitrogen MA1-20832 or CENP-T/W; 2.5mg/L; [13]; Vimentin; 1:500 Invitrogen PA1-16759 and Anti-Centromere Protein Antibody 1:100; Antibodiesinc; 15-234) and 0.4 U/µL RiboLock RNase inhibitor (Thermo Fisher Scientific; EO0384) diluted in 2.5% BSA in PBS for 1 hour at room temperature. Cells were washed three times with PBS-T and incubated with conjugated secondary antibodies (CENP-A screen: α-mouse AF488 Jackson ImmunoResearch; 715-545-150; CENP-T/ACA screen: α-rabbit AF750, Invitrogen A21039, α-human Cy2; Jackson ImmunoResearch; 709-225-149, α-chicken AF647; Invitrogen; A21449) diluted 1:500 and 0.4 U/µL RiboLock RNase inhibitor in 2.5% BSA in PBS for 45 minutes at room temperature in the dark. The reverse transcription reaction was set up as follows: 1X RevertAid RT buffer (Thermo Fisher Scientific; EP0452), 250 µM dNTPs (New England Biolabs; N0447L), 1 µM each biotinylated reverse transcription primer (oKC1015 /5Biosg/GTTGATAACGGACTAGCCTTATTTC), 200 µg/mL recombinant albumin (rAlbumin) (New England Biolabs; B9200S), 0.8 U/µL RiboLock RNase inhibitor, and 4.8 U/µL RevertAid H minus Reverse Transcriptase (Thermo Fisher Scientific EP0452). The cells were incubated in the reverse transcription reaction overnight at 37°C shaking at 135 rpm. Cells were washed five times with PBS and incubated in 20 µg/mL Streptavidin (New England Biolabs; N7021S) and 100 µg/mL rAlbumin (New England Biolabs; B9200S) in PBS for 15 minutes. Cells were washed three times with PBS-T and fixed with 3% formaldehyde and 0.1% glutaraldehyde in 1X PBS for 30 minutes. Following three PBS-T washes, the cells were incubated in gap-fill and ligation solution at 37°C for 5 min, followed by 45 °C for 90 minutes. The gap-fill and ligation solution was set up as follows: 1X Ampligase buffer (Lucigen; A3210K), 200 µg/mL rAlbumin (New England Biolabs; B9200S), 0.1 µM padlock probe (oDF59o /5Phos/GTTTCAGAGCTATGCTCTCCTGTTCGCCAAATTCTACCCACCACCCACTCTCCAaaggacgaaaCACCg), 50 nM dNTPs (New England Biolabs; N0447L), 0.02 U/µL TaqIT polymerase (Enzymatics; P7620L), 0.4 U/µL RNase H (Enzymatics; Y9220L), and 0.5 U/µL Ampligase (Lucigen; A3210K). Samples were then washed with PBS-T twice and incubated in rolling circle amplification (RCA) reaction solution overnight at 30°C. The RCA reaction solution was set up as follows: 1X Phi29 buffer (Thermo Fisher Scientific; EP0094), 250 µM dNTPs (New England Biolabs; N0447L), 200 µg/mL rAlbumin (New England Biolabs; B9200S), 5% glycerol, and 1 U/µL Phi29 DNA polymerase (Thermo Fisher Scientific; EP0094). Following RCA, cells were washed with PBS-T two times and again incubated with the same primary antibody solution minus RNase inhibitors for 45 minutes to enhance signal for the CENP-T screen. In both screens cells were again washed three times with PBS-T and incubated with the same secondary antibody solution for 45 minutes at room temperature in the dark. Following three PBS washes, wells were replaced with 2X SSC with 200 ng/mL DAPI and imaged for cellular phenotypes. Images were acquired using a Nikon Ti2 widefield microscope with 4 z-slices at 1.5 μm intervals. Following imaging, fluorophores were bleached using 1 mg/mL of lithium borohydride (LiBH_4_) [35] for 1 hour. Cells were washed three times with PBS three times and imaged for cellular phenotypes in 2X SSC with 200 ng/mL DAPI. Of note, other markers were also included in cellular phenotyping, but were not relevant to this study and are not reported here. Following imaging, fluorophores were again bleached in 1 mg/mL LiBH_4_ for 1 hour and then washed three times with 2X SSC.

Finally, *in situ* sequencing-by-synthesis was performed as described previously [18, 34]. First, 1 µM sequencing primer (oDF-663 anchP GCCAAATTCTACCCACCACCCACTCTCCAaaggacgaaaCACCg) in 2X SSC was hybridized for 30 minutes at 37°C, followed by two washes with PR2 (Illumina MS-103-1003). Cells were washed with PR2 and incubated in incorporation mix (MiSeq Nano kit v2 reagent 1, Illumina MS-103-1003) for 3 minutes at 60°C on a flat-top thermocycler. Cells were washed repeatedly with PR2, followed by three heated washes at 60°C for 5 minutes each. Prior to imaging of the first cycle of *in situ* sequencing, cells were incubated with FITC-conjugated anti-chicken secondary antibodies (diluted 1:500 in 2.5% BSA in PBS) for 45 minutes to label vimentin and aid in cellular segmentation. Cells were imaged in 2X SSC with 200 ng/mL DAPI. To proceed to the next cycle, samples were incubated in cleavage mix (Illumina kit reagent 4) for 6 minutes at 60°C. Samples were repeatedly washed with PR2, followed by two heated PR2 washes for 2 minutes at 60°C and one heated PR2 wash for 5 minutes at 60°C. Samples then returned to the incorporation step. Twelve cycles of *in situ* sequencing were performed. *In situ* sequencing images and cellular phenotypes were analyzed using Brieflow as previously described

#### Image analysis

All image analysis was performed with Brieflow v1.5.0 [21], with screen-specific parameters recorded in the accompanying analysis repositories (see Code availability). Both screens used HeLa cells expressing inducible Cas9 and a CROPseq-format sgRNA library targeting ∼5,000 fitness-conferring genes [18] (20,195 targeting sgRNAs, ∼4 per gene, plus 250 non-targeting control sgRNAs), imaged 78 hours after doxycycline induction. The CENP-A screen comprised three 6-well plates (18 wells); the CENP-T screen comprised five wells of a single 6-well plate. Each well was imaged as 377 *in situ* sequencing (SBS) tiles and 1,605 phenotype tiles. SBS and phenotype images were acquired at 10x and 20x respectively, giving a ∼4-fold combined magnification and binning difference between the two modalities (1,200 x 1,200 versus 2,400 x 2,400 pixel tiles).

#### Preprocessing

Raw ND2 images were converted to OME-Zarr and tile-level metadata (plate, well, tile, cycle or staining round) were extracted from file paths. Illumination correction fields were computed from a 10% random sample of tiles per plate–well combination and applied to all downstream images.

#### In situ sequencing

SBS images (DAPI plus four base channels G, T, A, C) were acquired over 11 cycles for the CENP-A screen and 12 cycles for the CENP-T screen, of which the first cycle was discarded because of poor quality, leaving 11 cycles in both screens. In the CENP-T screen, cycles 1 and 12 also carried a vimentin channel used for cell segmentation. Images were aligned across cycles by phase cross-correlation on a windowed center region with 2x subpixel upsampling; the CENP-A screen was aligned on the DAPI channel, and because the channel set differed between cycles in the CENP-T screen, alignment there used the maximum projection of the base channels. Aligned base channels were processed with a Laplacian-of-Gaussian filter (sigma = 1) followed by a maximum filter (width = 3 pixels). Candidate reads were identified from the standard deviation of filtered intensities across cycles by detecting local maxima within a 3-pixel (CENP-A) or 7-pixel (CENP-T) neighborhood, retaining peaks above a minimum intensity of 200 (CENP-A) or 100 (CENP-T). Nuclei and cells were segmented on the final cycle with Cellpose [36] (cyto3 model; nuclei diameter ∼10 pixels, cell diameter ∼19 pixels; flow threshold 0.4 for nuclei and 1.0 for cells, cell-probability threshold 0), using DAPI for nuclei and the C base channel (CENP-A screen) or vimentin (CENP-T screen) as the cytoplasmic channel; cells were retained only where they contained a nucleus. Base intensities at each peak were corrected for spectral cross-talk with a per-tile median-based linear transformation and called as the maximal channel. Reads were matched exactly to the 11-nucleotide sgRNA prefix in the library table (no error correction); each cell was assigned the barcode with the greatest number of supporting reads, and cells were retained for downstream analysis only if all mapped reads pointed to a single gene. Across wells, 60.6% (CENP-A) and 58.6% (CENP-T) of segmented SBS cells mapped to a single barcode (21.4 M and 5.2 M cells respectively).

#### Phenotype imaging and feature extraction

Nuclei and cell bodies were segmented with Cellpose (cyto3; CENP-A screen: nuclei diameter 43 pixels, cell diameter 76 pixels, WGA as the cytoplasmic channel; CENP-T screen: nuclei 40 pixels, cells 75 pixels, vimentin as the cytoplasmic channel; thresholds as above), and cells were reconciled to contain a nucleus. Morphological, intensity, texture, radial-distribution and pairwise colocalization features were extracted for each cell in nuclear, cellular and cytoplasmic compartments using Brieflow’s CellProfiler-emulating module [22] (1,661 features per cell in the CENP-A screen; 3,581 in the CENP-T screen). Foci features were additionally computed on the CENP-A and CENP-T channels respectively. Segmentation yielded 22.1 M (CENP-A) and 5.35 M (CENP-T) phenotype cells.

#### Spatial merging

SBS barcode assignments and phenotype feature profiles were linked at the single-cell level using Brieflow’s Delaunay-triangulation hashing [21]. Cell-centroid patterns from each modality were encoded as edge-length hash descriptors of Delaunay triangles and matched between tiles by nearest-neighbor search; six seed SBS tiles per well initialized the search, and further tile correspondences were discovered iteratively through RANSAC-based affine regression. Alignments were accepted if the affine determinant fell within 0.0563–0.0688 (0.9–1.1x the expected (1/4)^2^ scale factor) and the match score was at least 0.1. Individual cells were matched with a maximum coordinate distance of 2 pixels, and duplicate matches were resolved by nearest distance and then by mapping status. Merging recovered 85% (CENP-A) and 87% (CENP-T) of phenotype cells, of which 61% and 59% carried a single-gene barcode assignment.

#### Aggregation and statistical testing

Cells mapping to a single gene were retained. Features with more than 10% missing values were dropped, cells with more than 1% missing values were removed, and remaining values were imputed by k-nearest neighbors (k = 5). Intensity outliers (1%) were removed by local outlier factor on the per-channel mean intensities. Features were centered and scaled per batch (plate and well), projected by principal component analysis retaining 99% of variance, and batch-corrected by typical variation normalization: the PCA basis was re-fit on non-targeting control cells and each batch was whitened by its control covariance and re-colored with the pooled control covariance. Per-sgRNA and then per-gene profiles were computed as medians, and per-gene features were standardized as robust z-scores against non-targeting controls using the median absolute deviation. Gene profiles were built for DAPI+CENP-A, and DAPI+ACA+CENP-T from the two screens, respectively. Statistical significance was assessed by bootstrap resampling (100,000 iterations) on the full-panel profiles for a curated set of intensity (mean, integrated, edge, radial fraction, mass displacement), shape (area, solidity, form factor, eccentricity) and colocalization (Pearson correlation, Manders) features in the nuclear and cellular compartments (103 features in the CENP-A screen; 235 in the CENP-T screen). For each sgRNA, a null distribution of feature medians was built by drawing the sgRNA’s cell count with replacement from the cells of a randomly selected non-targeting sgRNA; gene-level nulls were the median across the gene’s sgRNAs; two-tailed empirical p-values were corrected by the Benjamini–Hochberg procedure across all gene–feature tests, and FDR < 0.05 was called significant.

#### Clustering

Genes represented by at least 5 cells were embedded by PHATE [37] (cosine distance, k = 10) and clustered with Leiden [38] at resolutions 6–10 (CENP-A) and 4–8 (CENP-T). Each clustering was benchmarked against CORUM complexes [39], KEGG pathways [40] and STRING interaction pairs [41] relative to a label-shuffled control. The figures use the profiles derived from DNA and CENP-A staining at resolution 9 (287 clusters) for the CENP-A screen, and the profiles derived from DNA, ACA and CENP-T staining at resolution 6 (248 clusters) for the CENP-T screen.

#### Single-feature readouts

CENP-A abundance was read out as the standardized mean nuclear CENP-A intensity. CENP-T abundance and centromeric recruitment were read out as the standardized mean nuclear CENP-T and ACA intensities and as the least-squares slope of CENP-T against ACA pixel intensity within the nucleus (ACA–CENP-T colocalization); bootstrap significance for volcano plots used the mean nuclear intensity and the nuclear DAPI–CENP-T Pearson correlation. Raw (unstandardized) per-gene means were computed directly from the single-cell data where shown.

## Code availability

Screen configuration, notebooks and figure scripts are in the damavand-analysis (CENP-A) and kazbek-analysis (CENP-T) repositories; Brieflow is available at github.com/cheeseman-lab/brieflow.

## Supplemental Figure Legends

**Supplemental Figure 1 (related to Figure 2):**
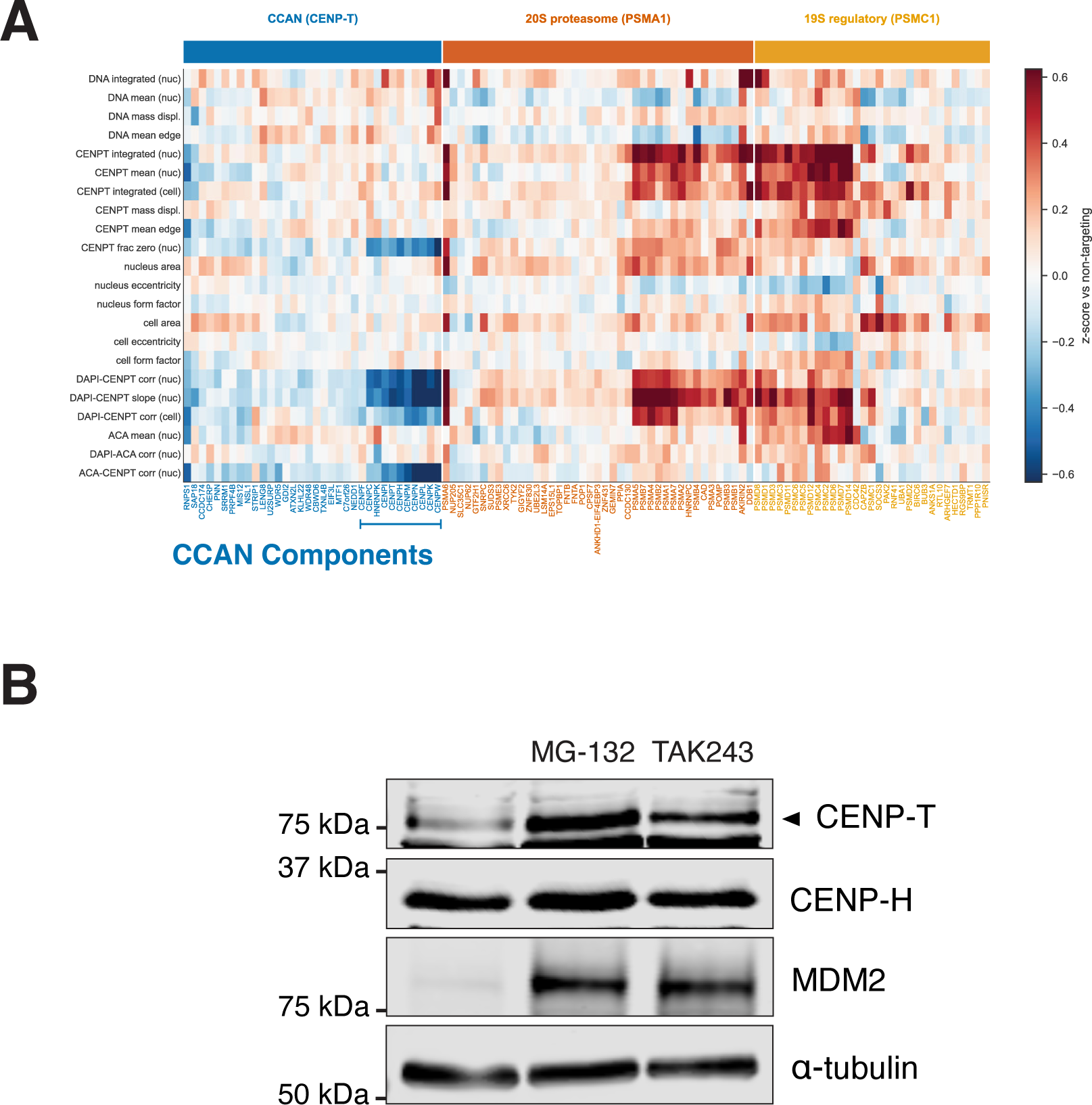
A. Heatmap showing representative features obtained from the OPS analysis of CENP-T localization for the centromere components and proteasome subunits. B. Western blots with anti-CENP-T or anti-CENP-H antibodies in proliferating cells and cells treated with MG-132 or TAK243. MDM2 serves as a control for protein stabilization and α-tubulin serves as a loading control.

**Supplemental Figure 2 (related to Figure 3):**
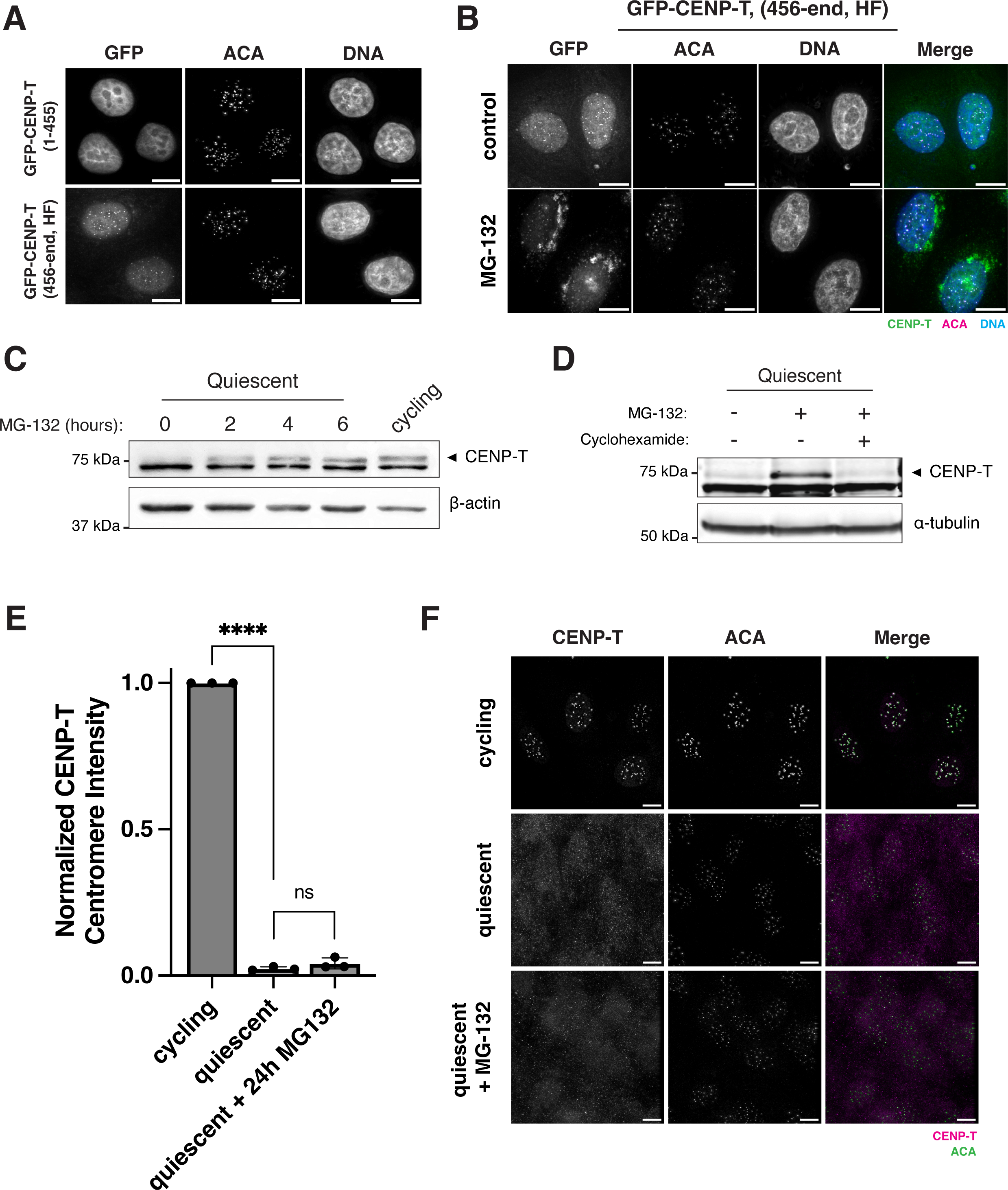
Quiescent cells degrade CENP-T. A. Representative immunofluorescence images of GFP-CENP-T cell lines. Top panels show an N-terminal CENP-T construct (amino acids 1-455). Bottom panels show a C-terminal histone fold CENP-T construct (amino acids 456-end). The N-terminal construct does not localize to the centromeres whereas the histone fold construct does. Cells were stained with GFP booster nanobody, anti-centromere antibody (ACA) and Hoechst for DNA. Scale bar, 10 µm. B. Representative immunofluorescence images of C-terminal histone fold CENP-T (amino acids 456-end) cell line with or without 24 hours of 10 µM MG-132 treatment. Cells were stained with GFP booster nanobody, anti-centromere antibody (ACA) and Hoechst for DNA. C. Western blot with anti-CENP-T antibodies of 7-day quiescent cells treated with 10 µM MG-132 for the indicated amount of time and of cycling cells. β-actin is used as a loading control. D. Western blot with anti-CENP-T antibodies of 7-day quiescent cells treated with 10 µM MG-132 and 50 µg/mL cycloheximide. α-tubulin serves as a loading control. E. Graph showing CENP-T centromere intensity as measure by immunofluorescence for cycling cells, quiescent cells, and quiescent cells treated with 10 µM MG-132 for 24 hours. Values were normalized to cycling condition. Bars represent mean ± standard deviation. **** represents p<0.0001. ns is not significant. Three biological replicates were conducted. 212, 447, and 425 centromeres were counted for cycling, quiescent, and quiescent with MG-132, respectively. F. Representative immunofluorescence images from E. Top panel: cycling cells, middle panel: quiescent cells in quiescence for 7 days, bottom panel: quiescent cells in quiescence for 7 days with 24 hours of 10 µM MG-132. Cells were stained with CENP-T antibody and anti-centromere antibody (ACA). Scale bar, 10 µm.

## References

1. McKinley, K.L., and Cheeseman, I.M. (2016). The molecular basis for centromere identity and function. Nat Rev Mol Cell Biol 17, 16–29.

2. Okada, M., Cheeseman, I.M., Hori, T., Okawa, K., McLeod, I.X., Yates, J.R., 3rd, Desai, A., and Fukagawa, T. (2006). The CENP-H-I complex is required for the efficient incorporation of newly synthesized CENP-A into centromeres. Nat Cell Biol 8, 446–457.

3. Foltz, D.R., Jansen, L.E.T., Black, B.E., Bailey, A.O., Yates, J.R., and Cleveland, D.W. (2006). The human CENP-A centromeric nucleosome-associated complex. Nat Cell Biol 8, 458–469.

4. Cheeseman, I.M., and Desai, A. (2008). Molecular Architecture of the Kinetochore-Microtubule Interface. Nat Rev Mol Cell Biol 9, 33–46.

5. Barnhart, M.C., Kuich, P.H., Stellfox, M.E., Ward, J.A., Bassett, E.A., Black, B.E., and Foltz, D.R. (2011). HJURP is a CENP-A chromatin assembly factor sufficient to form a functional de novo kinetochore. J Cell Biol 194, 229–243.

6. Jansen, L.E.T., Black, B.E., Foltz, D.R., and Cleveland, D.W. (2007). Propagation of centromeric chromatin requires exit from mitosis. J. Cell Biol. 176, 795–805.

7. McKinley, K.L., and Cheeseman, I.M. (2014). Polo-like Kinase 1 Licenses CENP-A Deposition at Centromeres. Cell 158, 397–411.

8. Foltz, D.R., Jansen, L.E., Bailey, A.O., Yates, J.R., 3rd, Bassett, E.A., Wood, S., Black, B.E., and Cleveland, D.W. (2009). Centromere-specific assembly of CENP-a nucleosomes is mediated by HJURP. Cell 137, 472–484.

9. Dunleavy, E.M., Roche, D., Tagami, H., Lacoste, N., Ray-Gallet, D., Nakamura, Y., Daigo, Y., Nakatani, Y., and Almouzni-Pettinotti, G. (2009). HJURP is a cell-cycle-dependent maintenance and deposition factor of CENP-A at centromeres. Cell 137, 485–497.

10. Fujita, Y., Hayashi, T., Kiyomitsu, T., Toyoda, Y., Kokubu, A., Obuse, C., and Yanagida, M. (2007). Priming of Centromere for CENP-A Recruitment by Human hMis18alpha, hMis18beta, and M18BP1. Dev Cell 12, 17–30.

11. Falk, S.J., Guo, L.Y., Sekulic, N., Smoak, E.M., Mani, T., Logsdon, G.A., Gupta, K., Jansen, L.E., Van Duyne, G.D., Vinogradov, S.A., et al. (2015). Chromosomes. CENP-C reshapes and stabilizes CENP-A nucleosomes at the centromere. Science 348, 699–703.

12. Hori, T., Amano, M., Suzuki, A., Backer, C.B., Welburn, J.P., Dong, Y., McEwen, B.F., Shang, W.H., Suzuki, E., Okawa, K., et al. (2008). CCAN makes multiple contacts with centromeric DNA to provide distinct pathways to the outer kinetochore. Cell 135, 1039–1052.

13. Gascoigne, K.E., Takeuchi, K., Suzuki, A., Hori, T., Fukagawa, T., and Cheeseman, I.M. (2011). Induced ectopic kinetochore assembly bypasses the requirement for CENP-A nucleosomes. Cell 145, 410–422.

14. Nishino, T., Rago, F., Hori, T., Tomii, K., Cheeseman, I.M., and Fukagawa, T. (2013). CENP-T provides a structural platform for outer kinetochore assembly. Embo J 32, 424–436.

15. Rago, F., Gascoigne, K.E., and Cheeseman, I.M. (2015). Distinct organization and regulation of the outer kinetochore KMN network downstream of CENP-C and CENP-T. Curr Biol 25, 671–677.

16. Nishino, T., Takeuchi, K., Gascoigne, K.E., Suzuki, A., Hori, T., Oyama, T., Morikawa, K., Cheeseman, I.M., and Fukagawa, T. (2012). CENP-T-W-S-X forms a unique centromeric chromatin structure with a histone-like fold. Cell 148, 487–501.

17. Sissoko, G.B., Tarasovetc, E.V., Marescal, O., Grishchuk, E.L., and Cheeseman, I.M. (2024). Higher-order protein assembly controls kinetochore formation. Nat Cell Biol 26, 45–56.

18. Funk, L., Su, K.C., Ly, J., Feldman, D., Singh, A., Moodie, B., Blainey, P.C., and Cheeseman, I.M. (2022). The phenotypic landscape of essential human genes. Cell 185, 4634–4653 e4622.

19. Kirby, B., Di Bernardo, M., Cheeseman, I.M., and Blainey, P.C. (2026). A one-week automated genome-wide optical pooled screen using OttoSeq. Genome Biol.

20. Bray, M.A., Singh, S., Han, H., Davis, C.T., Borgeson, B., Hartland, C., Kost-Alimova, M., Gustafsdottir, S.M., Gibson, C.C., and Carpenter, A.E. (2016). Cell Painting, a high-content image-based assay for morphological profiling using multiplexed fluorescent dyes. Nat Protoc 11, 1757–1774.

21. Di Bernardo, M., Kern, R.S., Cepeda Diaz, A.K., Mallar, A., Choi, S.J., Nutter-Upham, A., Lourido, S., Blainey, P.C., and Cheeseman, I. (2026). Brieflow: an integrated computational pipeline for high-throughput analysis of optical pooled screening data. Nat Commun 17.

22. McQuin, C., Goodman, A., Chernyshev, V., Kamentsky, L., Cimini, B.A., Karhohs, K.W., Doan, M., Ding, L., Rafelski, S.M., Thirstrup, D., et al. (2018). CellProfiler 3.0: Next-generation image processing for biology. PLoS Biol 16, e2005970.

23. McKinley, K.L., and Cheeseman, I.M. (2017). Large-Scale Analysis of CRISPR/Cas9 Cell-Cycle Knockouts Reveals the Diversity of p53-Dependent Responses to Cell-Cycle Defects. Dev Cell 40, 405–420.

24. McKinley, K.L., Sekulic, N., Guo, L.Y., Tsinman, T., Black, B.E., and Cheeseman, I.M. (2015). The CENP-LN complex forms a critical node in an integrated meshwork of interactions at the centromere-kinetochore interface. Mol Cell 60, 886–898.

25. Hayashi, T., Fujita, Y., Iwasaki, O., Adachi, Y., Takahashi, K., and Yanagida, M. (2004). Mis16 and Mis18 Are Required for CENP-A Loading and Histone Deacetylation at Centromeres. Cell 118, 715–729.

26. Shang, W.H., Hori, T., Westhorpe, F.G., Godek, K.M., Toyoda, A., Misu, S., Monma, N., Ikeo, K., Carroll, C.W., Takami, Y., et al. (2016). Acetylation of histone H4 lysine 5 and 12 is required for CENP-A deposition into centromeres. Nat Commun 7, 13465.

27. Khurana, S., Varma, D., and Foltz, D.R. (2024). Contribution of CENP-F to FOXM1-Mediated Discordant Centromere and Kinetochore Transcriptional Regulation. Mol Cell Biol 44, 209–225.

28. Marescal, O., and Cheeseman, I.M. (2020). Cellular Mechanisms and Regulation of Quiescence. Dev Cell 55, 259–271.

29. Swartz, S.Z., McKay, L.S., Su, K.C., Bury, L., Padeganeh, A., Maddox, P.S., Knouse, K.A., and Cheeseman, I.M. (2019). Quiescent Cells Actively Replenish CENP-A Nucleosomes to Maintain Centromere Identity and Proliferative Potential. Dev Cell 51, 35–48 e37.

30. Marescal, O., Su, K.C., Moodie, B., Taylor, N.J.L., and Cheeseman, I.M. (2026). The dynamics of centromere assembly and disassembly during quiescence. J Cell Biol 225.

31. Fulco, C.P., Munschauer, M., Anyoha, R., Munson, G., Grossman, S.R., Perez, E.M., Kane, M., Cleary, B., Lander, E.S., and Engreitz, J.M. (2016). Systematic mapping of functional enhancer-promoter connections with CRISPR interference. Science 354, 769–773.

32. van den Berg, J., A, G.M., Kielbassa, K., Feringa, F.M., Freire, R., and Medema, R.H. (2018). A limited number of double-strand DNA breaks is sufficient to delay cell cycle progression. Nucleic Acids Res 46, 10132–10144.

33. Stirling, D.R., Swain-Bowden, M.J., Lucas, A.M., Carpenter, A.E., Cimini, B.A., and Goodman, A. (2021). CellProfiler 4: improvements in speed, utility and usability. BMC Bioinformatics 22, 433.

34. Feldman, D., Singh, A., Schmid-Burgk, J.L., Carlson, R.J., Mezger, A., Garrity, A.J., Zhang, F., and Blainey, P.C. (2019). Optical Pooled Screens in Human Cells. Cell 179, 787–799 e717.

35. Radtke, A.J., Kandov, E., Lowekamp, B., Speranza, E., Chu, C.J., Gola, A., Thakur, N., Shih, R., Yao, L., Yaniv, Z.R., et al. (2020). IBEX: A versatile multiplex optical imaging approach for deep phenotyping and spatial analysis of cells in complex tissues. Proc Natl Acad Sci U S A 117, 33455–33465.

36. Stringer, C., and Pachitariu, M. (2025). Cellpose3: one-click image restoration for improved cellular segmentation. Nature methods 22, 592–599.

37. Moon, K.R., van Dijk, D., Wang, Z., Gigante, S., Burkhardt, D.B., Chen, W.S., Yim, K., Elzen, A.V.D., Hirn, M.J., Coifman, R.R., et al. (2019). Visualizing structure and transitions in high-dimensional biological data. Nat Biotechnol 37, 1482–1492.

38. Traag, V.A., Waltman, L., and van Eck, N.J. (2019). From Louvain to Leiden: guaranteeing well-connected communities. Sci Rep 9, 5233.

39. Giurgiu, M., Reinhard, J., Brauner, B., Dunger-Kaltenbach, I., Fobo, G., Frishman, G., Montrone, C., and Ruepp, A. (2019). CORUM: the comprehensive resource of mammalian protein complexes-2019. Nucleic Acids Res 47, D559–D563.

40. Kanehisa, M., Furumichi, M., Sato, Y., Matsuura, Y., and Ishiguro-Watanabe, M. (2025). KEGG: biological systems database as a model of the real world. Nucleic Acids Res 53, D672–D677.

41. Szklarczyk, D., Nastou, K., Koutrouli, M., Kirsch, R., Mehryary, F., Hachilif, R., Hu, D., Peluso, M.E., Huang, Q., Fang, T., et al. (2025). The STRING database in 2025: protein networks with directionality of regulation. Nucleic Acids Res 53, D730–D737.

